# Differential kin interactions between invasive and native plants: evidence from *Alternanthera philoxeroides* and its native congener

**DOI:** 10.64898/2026.03.03.709234

**Authors:** Yan Li, Ziying Tang, Xingliang Xu, Mark van Kleunen

**Affiliations:** College of Life Sciences, Nanjing Normal University, Nanjing, 210023, China; Key Laboratory of Ecosystem Network Observation and Modeling, Institute of Geographic Sciences and Natural Resources Research, Chinese Academy of Sciences, Beijing 100101 China; Ecology, Department of Biology, University of Konstanz, Universitätsstrasse 10, D-78457 Konstanz, Germany; Zhejiang Key Laboratory for Restoration of Damaged Coastal Ecosystems & Zhejiang Provincial Key Laboratory of Plant Evolutionary Ecology and Conservation, Taizhou University, Taizhou, Zhejiang, 318000, China

**Keywords:** alien plants, intraspecific interaction, kin selection hypothesis, niche partitioning theory, phenotypic plasticity, plant invasion

## Abstract

Reduced competition or facilitation between kin relative to nonkin can improve plant performance, particularly under resource-limited conditions. Understanding whether kin interactions differ between invasive and native species may provide insights into the mechanisms underlying the persistence and spread of invasive species, particularly for species that spread clonally. To explore this, we conducted a greenhouse experiment using the invasive *Alternanthera philoxeroides* and its native congener *A. sessilis* in China. For both species, we grew central plants without or with neighbors, and for the latter we had three intraspecific neighbor kinship treatments (kin only, nonkin only, and both kin and nonkin [mixed] neighbors). To test whether kinship effects are affected by resource limitation, we grew the plants under two watering conditions (well-watered and drought-stressed). Our findings revealed that at both the group (i.e., pot-level) and individual levels, invasive plants had a higher biomass production and experienced a less negative relative neighbor effect in kin groups than in nonkin groups, while these patterns were reversed in the native species. Although aboveground architecture of central plants did not differ significantly between kin and nonkin neighbors in either species, neighbor plants of the invasive species produced fewer nodes in kin groups than in nonkin groups, while the reverse was true for the native species. These patterns were not affected by the watering treatment. Together, these results indicate that while the native plants has stronger kin competition, the invasive species has reduced kin competition. Such reduced competition among kin in the invasive *Alternanthera philoxeroides* may enhance its population dominance and facilitate its spread.

## Introduction

Upon invasion by alien plants, the number of native species in a community is frequently reduced (Vilà, 2011; Thakur et al., 2025). Most studies exploring the mechanism underlying the dominance of invasive alien plants have focused on testing interspecific competition between invasive and native species. Many of these studies found that invasive plants often have higher competitive abilities than native plants (Kuebbing & Nunez, 2016; Golivets & Wallin, 2018). Although intraspecific competition is also a major determinant of the competitive outcome between species (Hart et al., 2018), few studies have examined the impact of intraspecific interactions on plant invasion dynamics. Because invasive plants typically form dense monoculture stands, intraspecific interactions may be more frequent in invasive species compared to native species (Williamson, 1996). Therefore, the strength of intraspecific interactions may be an important factor in the success of invasive species (Mangla et al., 2011; Zhang et al., 2019).

Many plants, especially those with limited dispersal mechanisms or clonal reproduction, tend to grow in proximity to relatives (Cheplick, 2022). Such kin interactions represent a primary form of intraspecific interactions that occur throughout a plant’s life. The kin selection hypothesis explains the outcome of kin interactions and suggests that plants are capable of kin recognition (i.e., the ability to distinguish genetically related individuals from strangers, Hamilton, 1964) and subsequently reduce competition with relatives while competing more intensely with unrelated individuals (Dudley et al., 2013; Mazal et al., 2023). For example, kin groups may exhibit enhanced growth or biomass production due to cooperative interactions (Ninkovic, 2003; Biedrzycki et al., 2010). Although there is support for the kin recognition hypothesis both in native and invasive species (Dudley & File, 2007; Murphy & Dudley, 2009; Crepy & Casal, 2015; Yang et al., 2018), few studies have tested whether the outcome of kin interactions differs between congeneric invasive and native species (see Lee et al., 2021; Zheng et al., 2021). If there are any differences, they could provide novel insights into the mechanisms underlying the dominance and spread of invasive species in their non-native ranges.

Kin interactions can shift from strong competition to reduced competition or facilitation under stressful conditions, such as drought or nutrient limitation (He et al., 2013; West et al., 2021). Specifically, Ploughe et al. (2019) proposed a framework describing drought-induced temporal shifts in plant–plant interactions and suggested that these interactions become more facilitative when drought stress increases but shift again to competition when stress is alleviated. In many regions, droughts are increasing in frequency because of global climate change and may have severe consequences for plant invasions (Sanders et al., 2025). Developing a more thorough understanding of kin interactions in invasive species under drought conditions is therefore critical for predicting the future spread of these species. Although drought stress often suppresses plant growth of native and invasive plants (Liu et al 2017), its potential to modify intraspecific interactions remains less explored. We therefore hypothesize that drought alters kin interaction dynamics, and that a greater change in these dynamics may confer an advantage to the invasive species under drought conditions.

*Alternanthera philoxeroides* (Mart.) Griseb. is an amphibious perennial herb native to South America. It was introduced to China in the 1930s and became a noxious invasive weed by the end of the last century. Invasive accessions of *A. philoxeroides* in China primarily rely on clonal reproduction and frequently form dense monocultures (Pan et al., 2007). This means that most ramets of this species are typically surrounded by kin. Furthermore, *A. philoxeroides* consistently dominates over its native congener *A. sessilis* (L.) R. Br. ex DC. when the two species co-occur (Lu et al., 2013, 2015; Gao et al., 2023). Therefore, *A. philoxeroides* and *A. sessilis* provide an ideal system for investigating differences in kin interactions between an invasive and a native species.

In this study, we conducted a greenhouse experiment to test whether the patterns of kin interaction differ between the invasive species *A. philoxeroides* and its native congener *A. sessilis* at both the group and individual levels. Since both species can reproduce clonally, we used ramets for propagation to ensure that kin are 100% genetically identical. This is important because variation in kin-interaction outcomes may be associated with the degree of genetic relatedness among kin originating from seed reproduction (Subrahmaniam et al., 2018; Mazal et al., 2023). Additionally, since the outcome of kin interactions may be altered by stress, we grew plants under both drought-stress and well-watered (i.e. control) conditions. We compared the performance of the native and the invasive species at both the group and individual levels. Specifically, we addressed the following questions: 1) Does the outcome of kin interactions differ between the native and the invasive species? 2) Is kin recognition and discrimination exhibited in these species3) How does water limitation modulate the kin interactions of the native and invasive species?

## Material and methods

### Study species

*Alternanthera philoxeroides* (Mart.) Griseb. (alligator weed, Amaranthaceae) is a perennial herb from South America that has become naturalized in many regions worldwide. Its stolons are long and branched, and at each node, it can produce roots and new shoots. These vegetative offspring (ramets) can become independent upon disconnection. In China, *A. sessilis* (L.) R. Br. ex DC. is a native congener of *A. philoxeroides*. While *A. philoxeroides* in China propagates solely through clonal reproduction, *A. sessilis* can propagate via seeds and stolon buds (Pan et al., 2007; Lu et al., 2015).

We selected four populations of the invasive species *A. philoxeroides* and four populations of the native species *A. sessilis* (kindly provided by Dr. Xinmin Lu, Table S1). Genetic variation in *A. philoxeroides* in China is low (Ye et al., 2003; Xu et al., 2003; Wang et al., 2024). Therefore, to increase the likelihood that we had collected different genotypes, the selected populations were at least 165 km apart for the invasive species, and 257 km for the native species. Before our experimental trials started, we vegetatively propagated the eight populations from stolon fragments at least three times to eliminate potential maternal environmental carry-over effects.

### Experimental setup

We conducted the experiments in a greenhouse of the Northeast Institute of Geography and Agroecology of the Chinese Academy of Sciences. On 14 July 2021, we chose healthy and strong stolons from a single maternal plant from each population. These stolons were then cut into single-node fragments, and approximately 120–150 of such fragments were planted in trays. The cultivation substrate was potting soil (Pindstrup Plus, Pindstrup Mosebrug A/S, Denmark; pH: 6; 120.0 mg L^-1^ N; 12.0 mg L^-1^ P; 400.0 mg L^-1^ K; 28.0 mg L^-1^ Mg; 0.4 mg L^-1^ B; 2.0 mg L^-1^ Mo; 1.7 mg L^-1^ Cu; 2.9 mg L^-1^ Mn; 0.9 mg L^-1^ Zn; 8.4 mg L^-1^ Fe). The trays were randomly assigned to positions on a greenhouse bench. The temperature in the greenhouse ranged between 22 °C and 28 °C, and natural lighting was approximately 75% of the outdoor light levels.

Seven days after planting (21 July 2021), we started to transplant plantlets to rectangular 6-L pots (length × width × height: 26 cm × 16 cm × 14 cm) filled with soil collected from an abandoned agricultural field. This soil was nutrient-rich and thus posed no nutrient limitation to plant growth. We planted one plantlet (central plant) in the center of each pot and two neighbor plantlets on either the left or right side of the central plant (Figure 1). The distance from each neighbor to the central plant was 6.5 cm. Our experiment design was as follows. The central plants were designed to grow with three different neighbor kinship configurations: (1) kin only (the central plant grew with two kin neighbors); (2) nonkin only (the central plant grew with two different nonkin neighbors) and (3) mixed (the central plant grew with one kin and one nonkin neighbor). Kin neighbors were plantlets from the same maternal plant as the central plants (i.e. they were genetically identical), whereas nonkin neighbors were plantlets from different maternal plants of the same species as the central plants. In addition, to assess overall neighbor effects on the central plants, we also grew central plants alone in pots as controls (see Figure 1a for experimental design). Considering that intraspecific interactions could be affected by drought stress, we applied two watering treatments (well-watered and drought).

**Figure 1.**
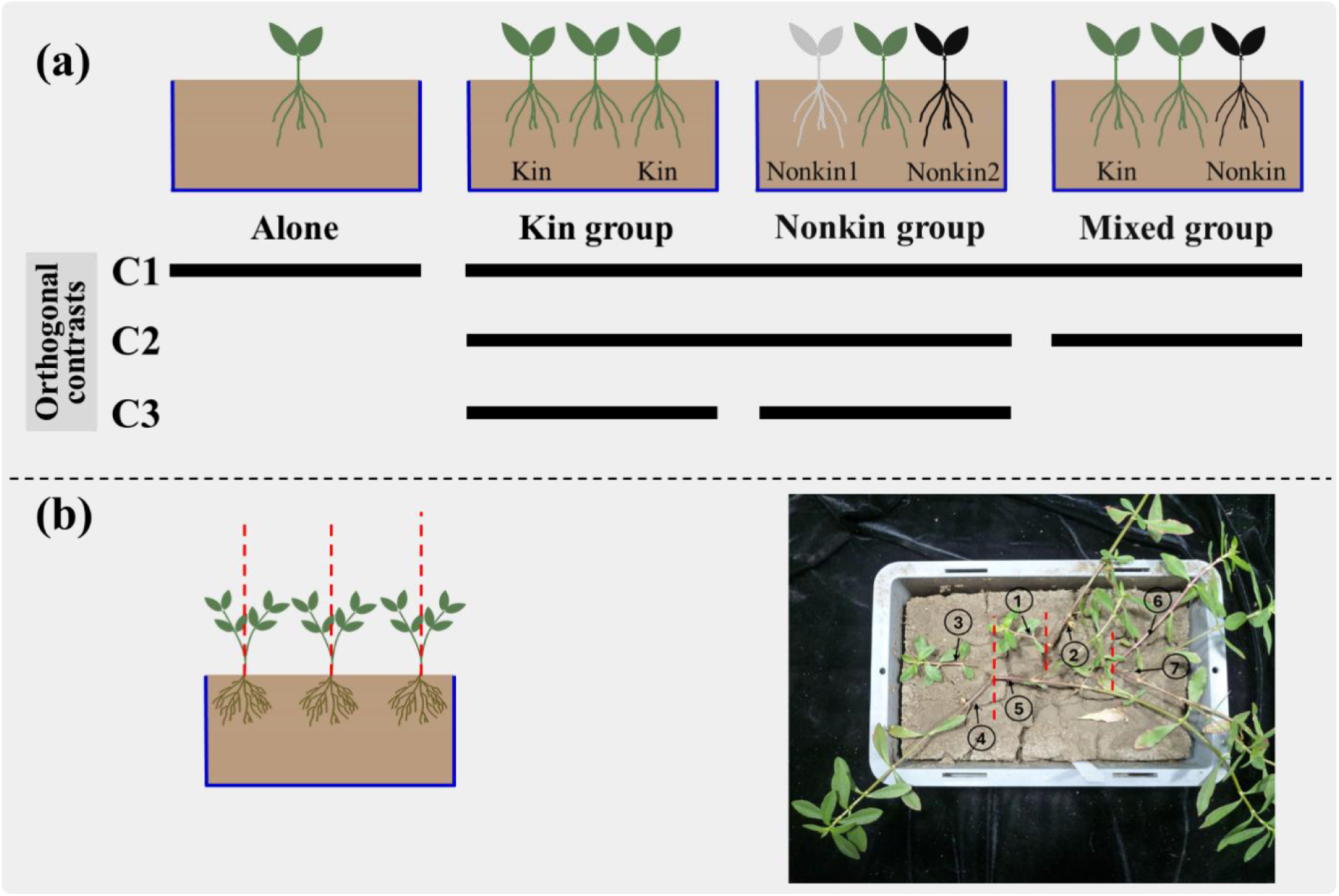
Schematic diagram of the experimental design (a) and the separation of the two aboveground halves of each plant (b). The plant in the middle of each pot is the central plant, and it is either grown alone or with two neighbor plants. Plants that have the same color are kin, and plants of different colors are nonkin. The central plants was grown either with two kin neighbors (kin group), two different nonkin neighbors (nonkin group) or one kin and one nonkin neighbor (mixed group). The four different neighbor treatments were coded as dummy variables to have three specific non-orthogonal contrasts (C1, C2, and C3) (a). To analyze whether plants grew towards kin or nonkin, we divided each plant into two parts along a vertical line (the red dotted line) through the center of each plant. In the example shown in the photo, the left and right sides of the central plant consist of branches ① and ②, respectively; the left side of the left neighbor consist of branches ③ and ④, and its right side consists of branch ⑤; for the right neighbor, the right side consists of branches ⑥ and ⑦, and there are no branches on its left side (b).

Based on this experimental design, we selected 60 similar-sized plantlets from each maternal plant for transplanting. Twenty-four plantlets (4 neighbor types [without neighbor, kin neighbors, mixed neighbors, and nonkin neighbors] × 2 watering treatments × 3 replications) from each maternal plant were selected to serve as central plants, and the remaining 36 plantlets served as neighbors ([3 plantlets as kin neighbors + 3 plantlets as nonkin neighbors] × 2 watering treatments × 3 replicates). This design resulted in 192 pots (8 maternal plants × 4 neighbor types × 2 watering treatments × 3 replicates), and a total of 480 plants. All pots were randomly assigned to positions on four benches in the greenhouse.

We watered all pots to saturation at transplanting. Subsequently, we provided sufficient water to keep the substrate moist throughout the entire experiment for the pots allocated to the wet treatment, which served as the control. As wilting is a reliable universal indicator of drought stress (Huai et al., 2003), we withheld water from plants in the drought treatment until plant wilting was observed. Therefore, we daily checked all plants in the drought treatment for signs of wilting, and supplied 100 ml (from 9 to 28 August, in the early stage of plant growth) or 300 ml (from 29 August to 11 September, in the mature stage of plant growth) of water to each pot with at least one wilting plant.

### Measurements

On 11 September 2021 (59 days after the start of the experiment), we concluded the experiment. To test whether neighbor kinship affects the distribution of aboveground architecture in plants, we divided each plant into two parts along a vertical line through the planting point (Figure 1b). We then measured the stolon length and counted the nodes and branches on each side (left and right) of each plant within the pots. Subsequently, we harvested the belowground and aboveground biomass in each pot. For the aboveground parts, we harvested each plant per pot separately. For the belowground parts, all roots per pot were harvested together because of the difficulty of assigning roots to individual plants when grown with neighbors. All plant material was dried at 70°C for at least three days and then weighed on a scale with an accuracy of 0.0001 g.

The measured growth traits are ecologically relevant as they are related to plant fitness and competitive ability. Stolon length and node number are associated with vegetative propagation because longer stolon and more nodes enable rapid spatial expansion and the formation of new ramets, thereby facilitating population spread and persistence (Oborny & Kun, 2001). The number of branches indicates the plant’s ability to produce new growth points, enhance light capture, and realize spatial dominance in dense stands (Pearcy et al., 2005). Biomass serves as an integrative indicator of fitness that reflects the plant’s capacity to acquire and allocate resources for survival and reproduction (Younginger et al., 2017).

### Statistical analyses

To assess the neighbor effect on the central plants, we calculated the relative neighbor effect (RNE) of central plants using the following formula (Markham & Chanway, 1996):

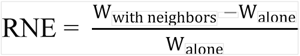

Where W_with neighbors_ and W_alone_ are the values of aboveground biomass of the central plant when grown with neighbors and grown alone. An index value around 0 indicates that neighbors had no effect on the central plant. A value larger than 0 indicates that neighbors had a positive effect, while a value smaller than 0 indicates that neighbors had a negative effect.

Moreover, to assess whether kinship of the neighbors influences the spatial distribution of aboveground architecture, we calculated the log response ratio (LRR) for three traits: stolon length, node number, and branch number. The LRR was defined as the natural logarithm of the ratio of allocation towards the kin neighbor to that towards the nonkin neighbor, using the following formula (Hedges et al., 1999):

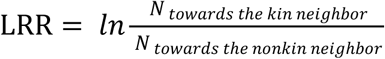

where N_towards the kin neighbor_ represents the values of stolon length, node number or branch number of the central plants towards the kin neighbor; and N_towards the nonkin neighbors_ represents the values of these traits of the central plants towards the nonkin neighbor. For central plants grown alone, with two kin neighbors, or with two nonkin neighbors (where no true kin–nonkin contrast existed), we randomly allocated their branches to dummy ‘kin’ and ‘nonkin’ neighbor categories for LRR calculation. LRR values are symmetrical around zero: positive values indicate that the central plant allocated more resources to the kin neighbor side, while negative values indicate greater allocation to the nonkin neighbor side.

To analyze the effects of neighbor type and its interactions with species status and watering treatment on plant performance, we used linear and generalized linear mixed effects models implemented in the *lme* function of the ‘nlme’ package (Pinheiro et al. 2020) and the glmer function of the ‘lme4’ package (Bates et al. 2015) in R 4.2.3 (R Core Team 2023).The fixed part of models always included species status (native and invasive), neighbor type (without neighbor, kin neighbors, mixed neighbors, and nonkin neighbors; see Figure 1a), watering treatment (wet and drought), and their interactions. The random part of models always included the maternal population of the central plant.

For biomass production and the RNE, we used Gaussian error distributions. To improve normality and homogeneity of the residuals, we used natural log-transformation prior to analysis. For node number and branch number, we used Poisson error distributions.

We coded the four neighbor types as three dummy variables (C1, C2, and C3, Table S2; see Schielzeth [2010] for an explanation of dummy variables). The dummy variable C1 tests whether the presence of neighbors had a significant effect on plant performance by comparing plants grown alone to those grown with neighbors (averaged across kin, mixed, and nonkin neighbor types). The dummy variable C2 tests whether the kin effect is additive by comparing plants grown with mixed neighbors (one kin and one nonkin neighbors) to the average of those grown with two kin neighbors and those grown with two nonkin neighbores. The dummy variable C3 tests whether neighbor kinship (kin vs. nonkin) influence plant performance by comparing plants grown with two kin neighbors to those grown with two nonkin neighbors (Figure 1a).

We tested the significance of the fixed effects (including the dummy variables) and all interactions by removing them from the model in hierarchical order, starting with the highest order interactions, and performing model comparisons using likelihood-ratio tests (Zuur et al., 2009).

## Results

### Biomass production per pot

The native species produced significantly more total biomass (aboveground biomass + root biomass) and root biomass per pot than the invasive alien species (total biomass: 18.02 g vs.8.43 g; root biomass: 2.86 g vs. 1.53 g; significant effect of Status in Table 1; Figure 2a, b). Drought resulted in a reduction of biomass production relative to the control (total biomass: 21.25 g to 5.20 g; root biomass: 3.40 g to 0.99 g; significant effect of Watering treatment in Table 1; Figure 2a, b).

**Figure 2.**
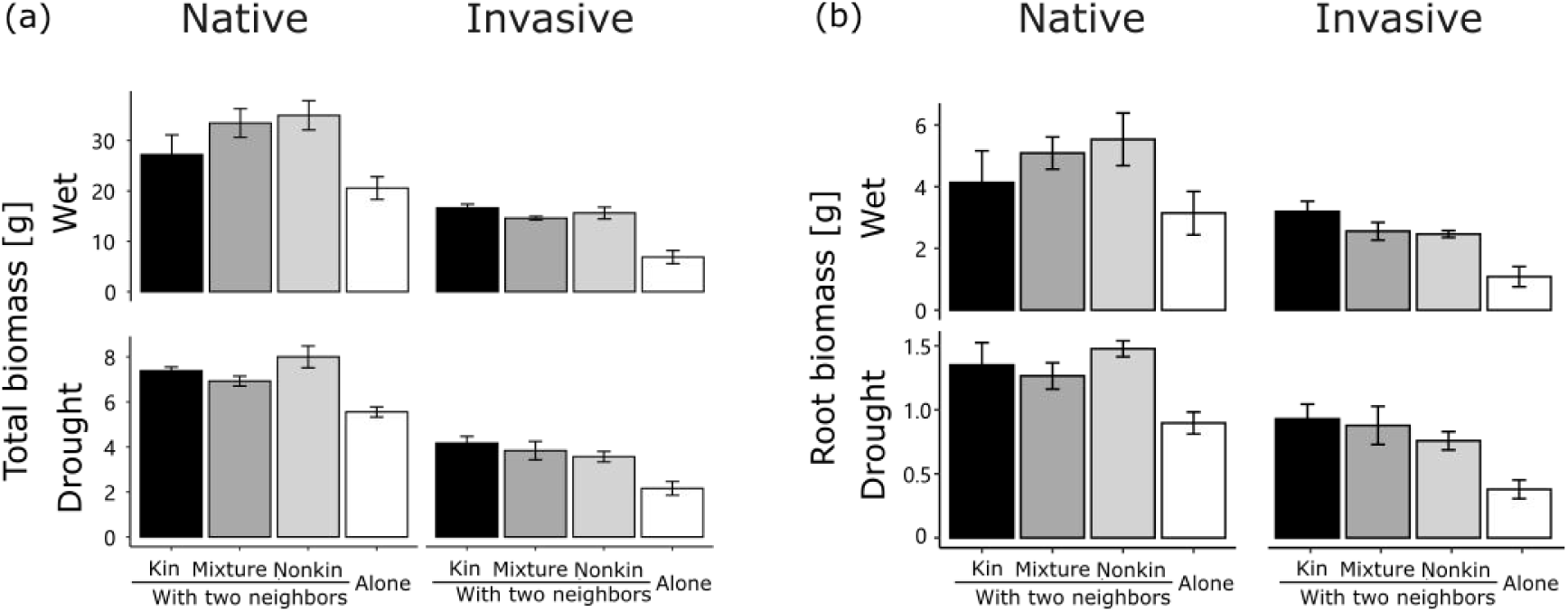
Total biomass (a) and root biomass (b) per pot when central plants were grown with kin neighbors (black), with mixed neighbors (dark grey), with nonkin neighbor (light grey) or alone (white) for native and invasive plants in the wet and drought treatments. Error bars are standard errors of the mean of population-means (n = 4 populations; each population mean was based on 3 individual plants).

**Table 1.**
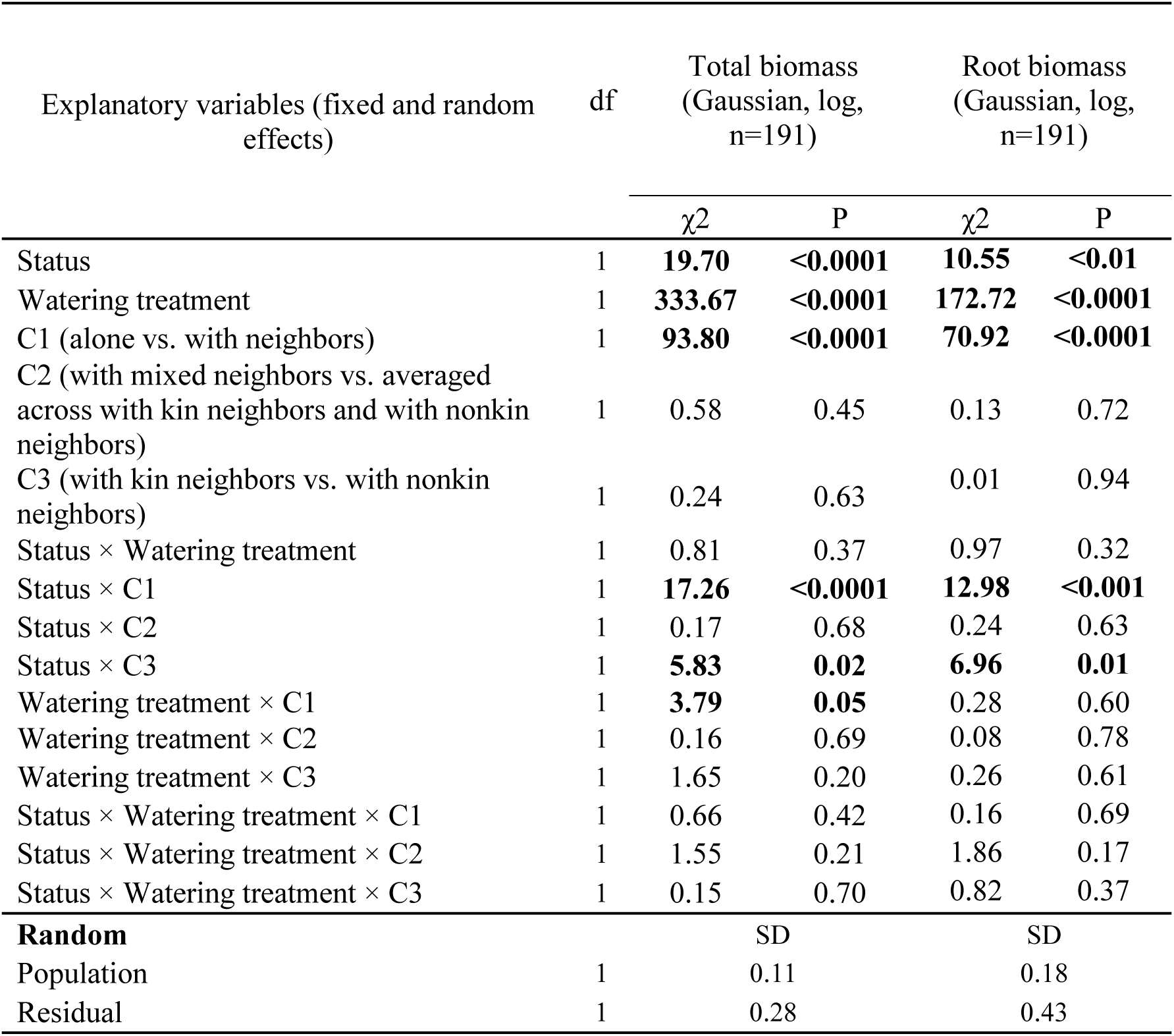
Results of linear mixed-effect models used to test the effects of species status (invasive vs. native), watering treatment (wet vs. drought), neighbor type (coded as dummy variables: C1, C2, C3; dummy variables were coded to allow testing of the specific contrasts listed in Figure 1) and all their interactions on aboveground biomass and root biomass per group. Model error distribution, applied transformations, and sample size are indicated in brackets. Significant effects (P < 0.05) are in bold.

Not surprisingly, pots with central plants and neighbors produced more total biomass than those with central plants only (total biomass:14.70 g vs. 8.79 g; root biomass:2.47 g vs. 1.38 g; significant effects of C1 in Table 1; Figure 2a, b). However, this effect was stronger for the invasive species than for the native species (significant Status × C1 interaction in Table 1). For the pots with neighbors, the invasive species produced more biomass when the neighbors were kin of the central plant than when they were nonkin (with 10.40 g vs. 9.59 g for total biomass, and 2.06 g vs. 1.61 g for root biomass), whereas the opposite was true for the native species (with 17.30 g vs. 21.50 g for total biomass, and 2.74 g vs. 3.51 g for root biomass; significant Status × C3 interaction in Table 1; Figure 2a, b). No significant difference was found between mixed groups (i.e. pots with kin and nonkin neighbors) and the average of pots with two kin neighbors and the pots with two nonkin neighbors (no significant effect of C2 in Table 1), suggesting that the effects of kin neighbors were additive.

### Neighbor effects on aboveground biomass and stolon growth of central plants

The presence of neighbors had a significant negative effect on the biomass production of central plants, with the relative neighbor effect (RNE) ranging from −0.14 to −0.66 (Figure 3). The RNE was significantly more negative for the native species than for the invasive species (−0.56 vs. −0.35; Table 2; Figure 3). Furthermore, for both species, drought increased the magnitude of the negative RNE relative to the control (−0.53 to −0.37; Table 2; Figure 3).

**Figure 3.**
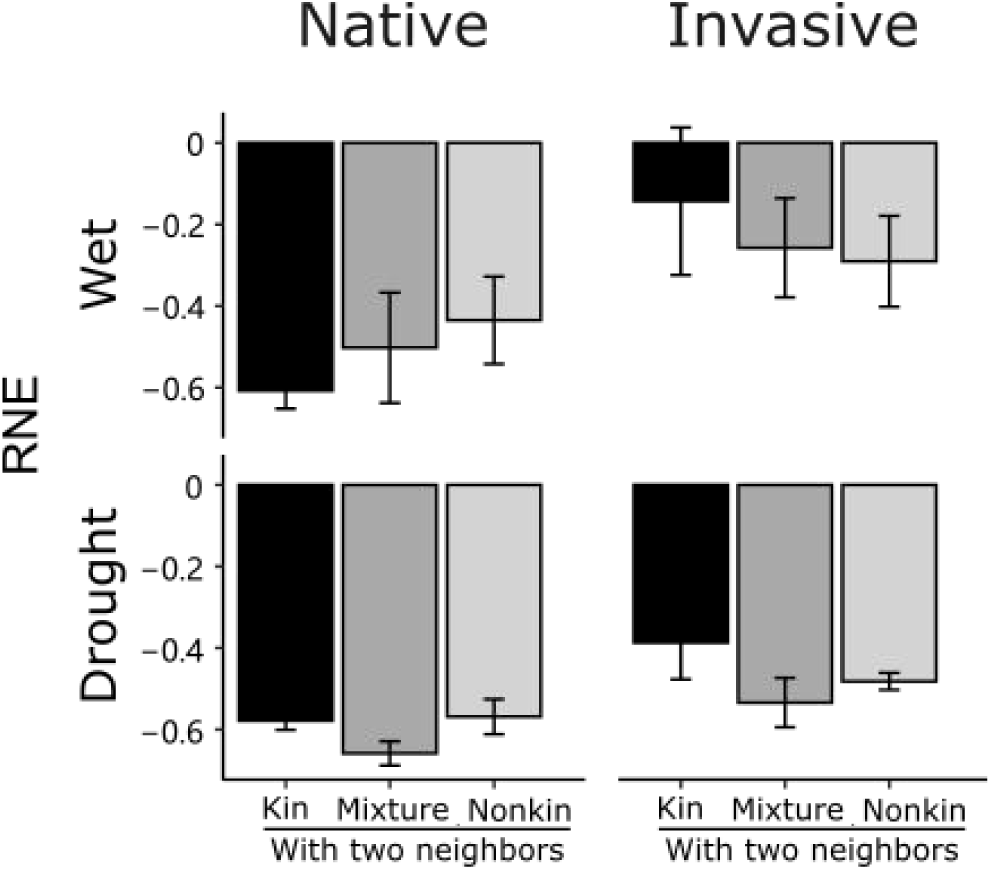
The relative neighbor effect (RNE) of aboveground biomass of central plants for native and invasive species under wet and drought conditions. Plants were grown in kin groups (black), mixed groups (dark grey) or nonkin groups (light grey). Error bars are standard errors of the mean of population-means (n = 4 populations; each population mean was based on 3 individual plants).

**Table 2.**
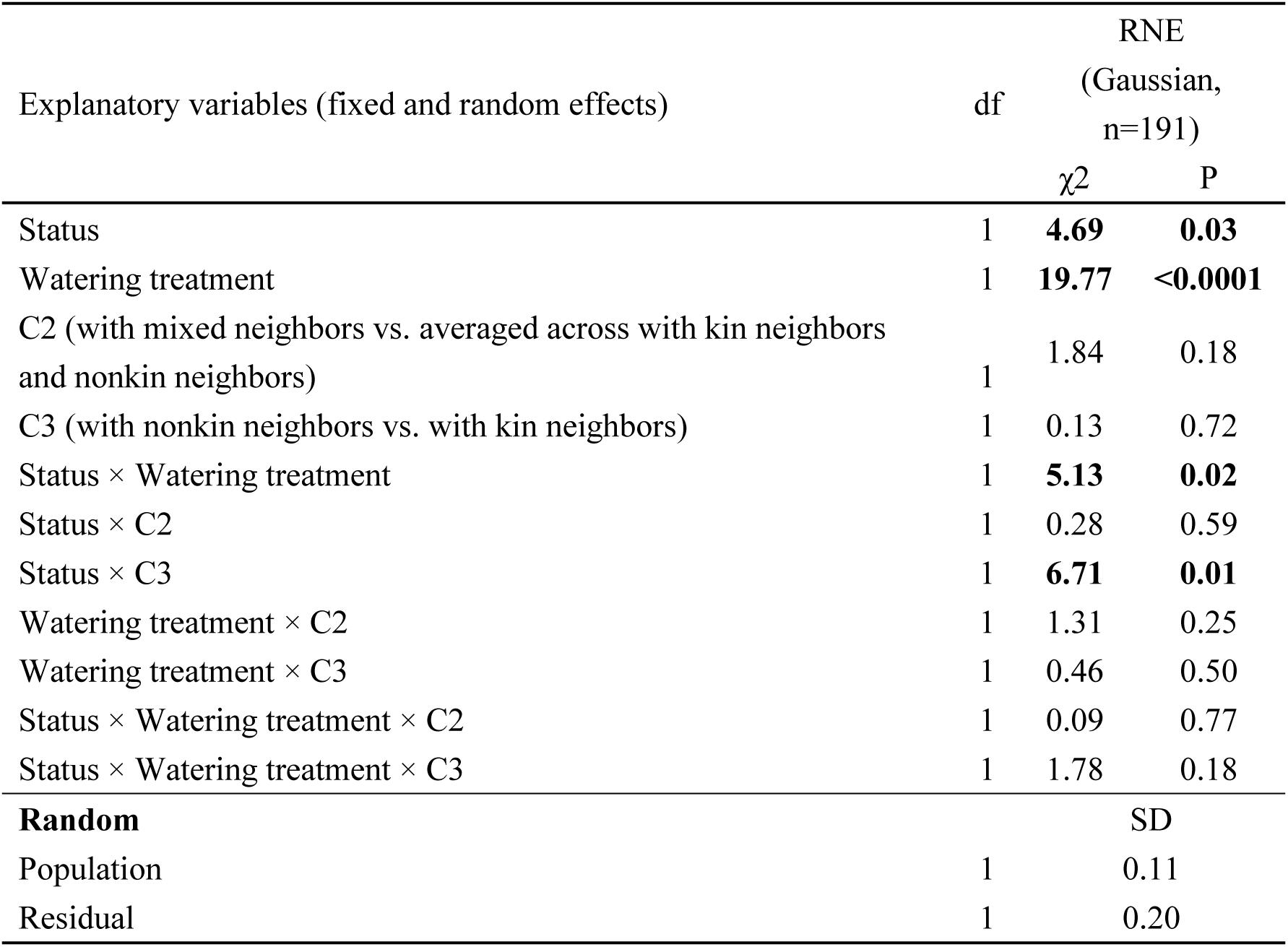
Results of linear mixed-effect models used to test the effects of species status (invasive vs. native), watering treatment (wet vs. drought), neighbor type (coded as dummy variables: C2, C3; dummy variables were coded to allow testing of the specific contrasts listed in Figure 1) and all their interactions on the relative neighbor effect (RNE) of aboveground biomass of central plants. Model error distribution and sample size are indicated in brackets. Significant effects (P < 0.05) are in bold.

| Explanatory variables (fixed and random effects) | df | RNE<br>(Gaussian,<br>n=191) |  |
| --- | --- | --- | --- |
| | | $\chi^2$ | P |
| Status | 1 | **4.69** | **0.03** |
| Watering treatment | 1 | **19.77** | **<0.0001** |
| C2 (with mixed neighbors vs. averaged across with kin neighbors and nonkin neighbors) | 1 | 1.84 | 0.18 |
| C3 (with nonkin neighbors vs. with kin neighbors) | 1 | 0.13 | 0.72 |
| Status × Watering treatment | 1 | **5.13** | **0.02** |
| Status × C2 | 1 | 0.28 | 0.59 |
| Status × C3 | 1 | **6.71** | **0.01** |
| Watering treatment × C2 | 1 | 1.31 | 0.25 |
| Watering treatment × C3 | 1 | 0.46 | 0.50 |
| Status × Watering treatment × C2 | 1 | 0.09 | 0.77 |
| Status × Watering treatment × C3 | 1 | 1.78 | 0.18 |
| **Random** |  | SD |  |
| Population | 1 | 0.11 |  |
| Residual | 1 | 0.20 |  |

For the invasive species, the RNE was less negative (indicating weaker competition) when the central plant had two kin neighbors than when it had two nonkin neighbors (−0.26 vs. −0.39), whereas the opposite tended to be true for the native species (−0.59 vs. −0.54). When we analysed the aboveground biomass of the central plant, we found the same pattern as for the RNE: central plants of the invasive species produced more biomass when they had kin neighbors (2.82 g vs. 2.39 g), while central plants of the native species produced less biomass when they had kin neighbors (4.41 g vs. 5.74 g; Table S3, Figure S1).

The central plants that had both one kin and one nonkin neighbour (i.e. mixed groups) allocated 48% of total stolon length toward the kin side and 52% toward the non-kin side. In line with this, the average log response ratio (LRR) was close to zero (0.02), and there were no significant effects of species status, watering treatment, neighbor kinship or any of their interactions (Table 3; Figure 4). Similarly, these factors did not significantly affect the LRR of node number and branch number (Table S4).

**Figure 4.**
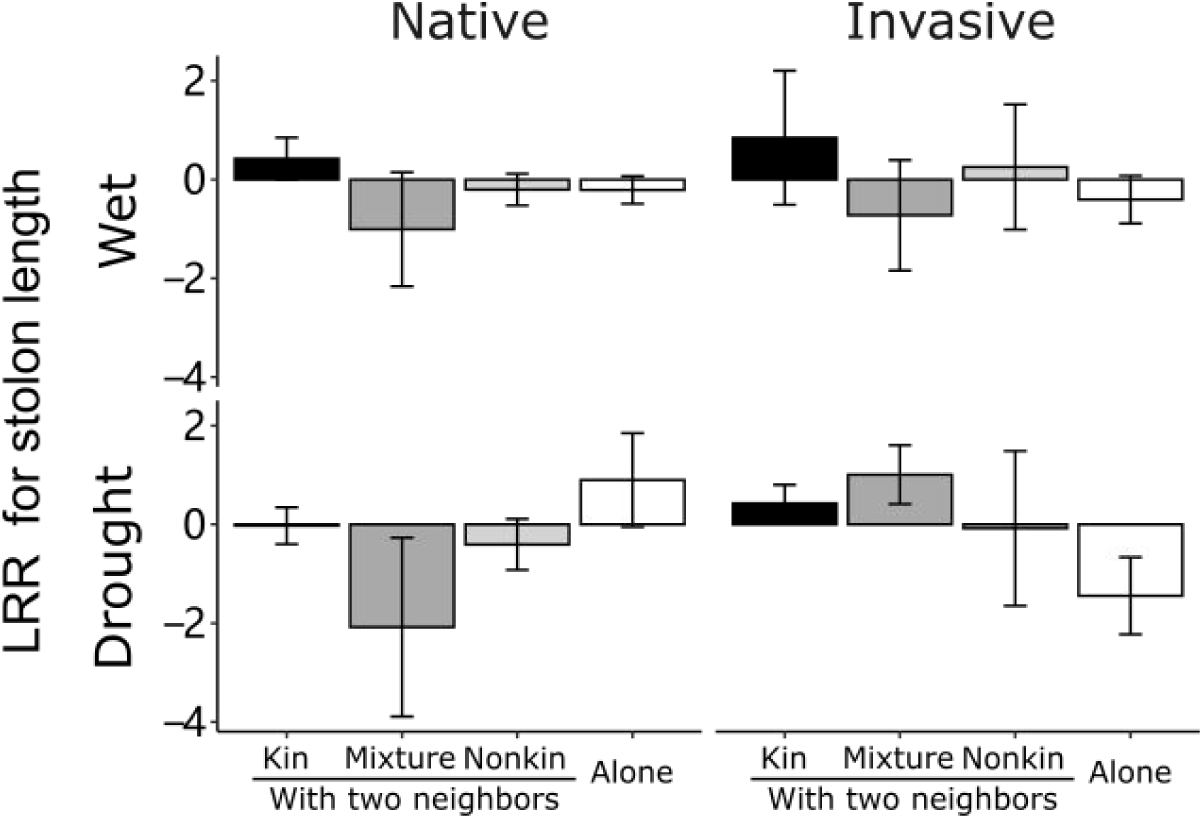
The log response ratio (LRR) for stolon length of central plants for the native and invasive species under wet and drought conditions. Plants were grown in kin groups (black), mixed groups (dark grey), nonkin groups (light grey), or alone (white). Error bars are standard errors of the mean of population-means (n = 4 populations; each population mean was based on 3 individual plants).

**Table 3.** Results of linear mixed-effect models used to test the effects of species status (invasive vs. native), watering treatment (wet vs. drought), neighbor type (coded as dummy variables:C1 C2, C3; dummy variables were coded to allow testing of the specific contrasts listed in Figure 1) and all their interactions on the log response ratio (LRR) of stolon length of central plants. Model error distribution and sample size are indicated in brackets. Significant effects (P < 0.05) are in bold.

| Explanatory variables (fixed and random effects) | df | LRR for stolon length<br>(Gaussian, n=190) |  |
| --- | --- | --- | --- |
| | | $\chi^2$ | P |
| Status | 1 | 0.46 | 0.50 |
| Watering treatment | 1 | 0.06 | 0.81 |
| C2 (with mixed neighbors vs. averaged across with kin neighbors and nonkin neighbors) | 1 | 0.11 | 0.74 |
| C3 (with kin neighbors vs. with nonkin neighbors) | 1 | 2.70 | 0.10 |
| Status × Watering treatment | 1 | 0.68 | 0.41 |
| Status × C2 | 1 | 0.02 | 0.89 |
| Status × C3 | 1 | **4.49** | **0.03** |
| Watering treatment × C2 | 1 | 1.41 | 0.23 |
| Watering treatment × C3 | 1 | 0.00 | 1.00 |
| Status × Watering treatment × C2 | 1 | 0.03 | 0.86 |
| Status × Watering treatment × C3 | 1 | 0.47 | 0.49 |
| **Random** |  | 0.01 | 0.92 |
| Population | 1 | 2.33 | 0.13 |
| Residual | 1 | 2.07 | 0.15 |

### Aboveground biomass production and branching of neighbor plants

Consistent with the pattern we found for the central plants, the average aboveground biomass of the two neighbor plants was significantly higher for the native than the alien species and was reduced by drought (Table S3, Fig. S1). Moreover, native neighbor plants produced less biomass in when they were in the kin-only treatment than when they were in the nonkin-only treatment (10.15 g vs. 12.26 g), whereas invasive neighbor plants showed a slight opposite trend (5.59 g vs. 5.51 g) (marginally non-significant Status × C3 interaction in Table S3; Fig. S1). However, the branching patterns contrasted with the biomass patterns. Native neighbor plants produced more branches in the kin-only treatment than in the nonkin-only treatment (45.42 vs. 36.21), while invasive neighbor plants showed the reverse trend (18.81 vs. 19.98) (Table 4; Figure 5).

**Figure 5.**
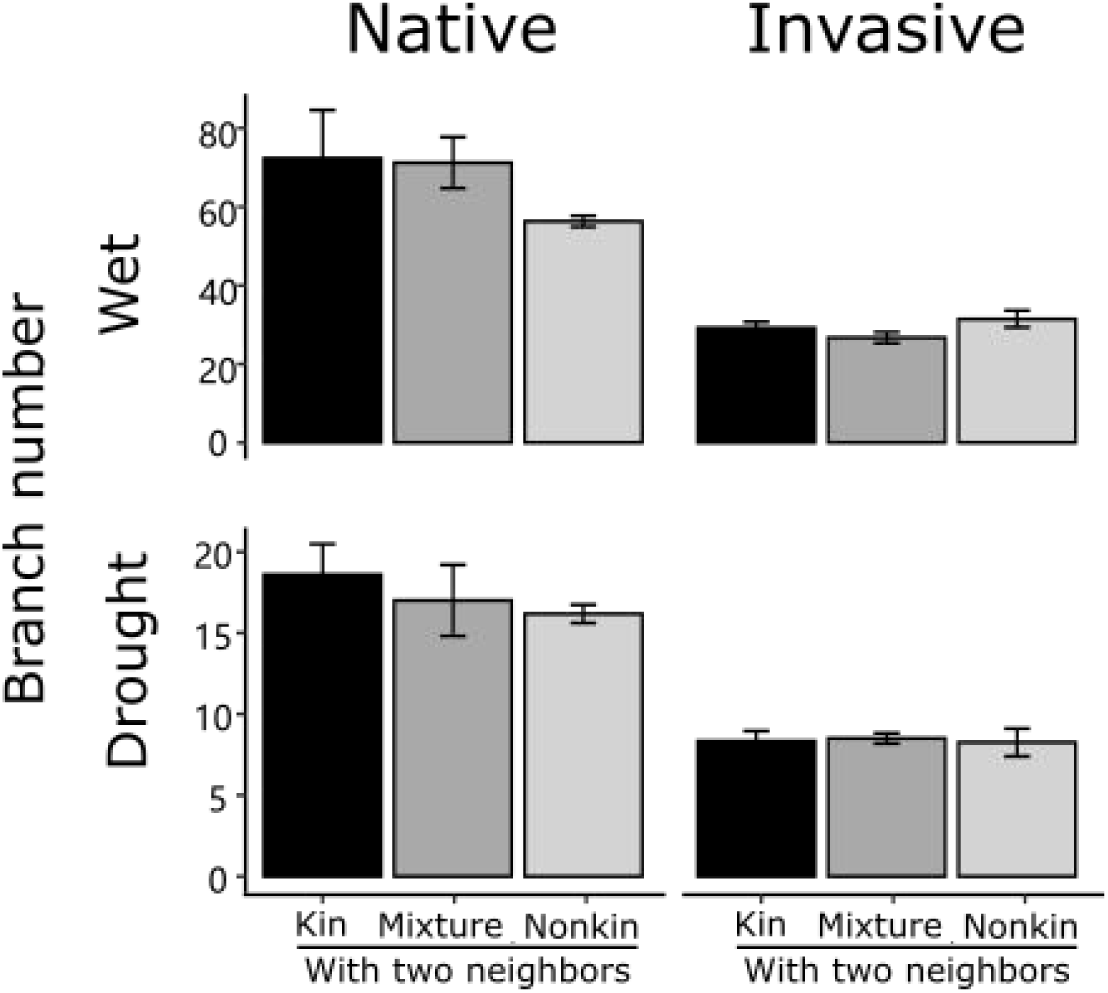
The branch number of the two neighbor plants for native and invasive species under wet and drought conditions. Plants were grown in kin groups (black), mixed groups (dark grey) or nonkin groups (light grey). Error bars are standard errors of the mean of population-means (n = 4 populations; each population mean was based on 3 pots, each with two neighbor plants).

**Table 4.** Results of linear mixed-effect models used to test the effects of species status (invasive vs. native), watering treatment (wet vs. drought), neighbor type (coded as dummy variables: C1, C2, C3; dummy variables were coded to allow testing of the specific contrasts listed in Figure 1) and all their interactions on the branch number of two neighbor plants. Model error distribution, applied transformations, and sample size are indicated in brackets. Significant effects (P < 0.05) are in bold.

| Explanatory variables (fixed and random effects) | df | Branch number<br>(poisson,<br>n=143) |  |
| --- | --- | --- | --- |
| | | $\chi^2$ | P |
| Status | 1 | **23.41** | **<0.0001** |
| Watering treatment | 1 | **265.89** | **<0.0001** |
| C2 (with mixed neighbors vs. averaged across with kin neighbors and nonkin neighbors) | 1 | 0.00 | 0.96 |
| C3 (with kin neighbors vs. with nonkin neighbors) | 1 | 3.31 | 0.07 |
| Status × Watering treatment | 1 | 1.08 | 0.30 |
| Status × C2 | 1 | 2.13 | 0.14 |
| Status × C3 | 1 | **4.25** | **0.04** |
| Watering treatment × C2 | 1 | 0.09 | 0.77 |
| Watering treatment × C3 | 1 | 0.03 | 0.87 |
| Status × Watering treatment × C2 | 1 | 1.52 | 0.22 |
| Status × Watering treatment × C3 | 1 | 0.69 | 0.41 |
| **Random** |  | SD |  |
| Population | 1 | 0.07 |  |
| Residual | 1 | 0.17 |  |

## Discussion

This study investigated intraspecific interactions with kin and nonkin individuals in the invasive species *A. philoxeroides* and its native congener *A. sessilis*. Our results demonstrate that kinship of the plants in each pot differentially influenced biomass production and aboveground architecture of the two species. Specifically, for the invasive species, the central plant experienced less competition by kin neighbors than by nonkin neighbors, and the biomass production per pot was increased when all the plants were kin, whereas the native *A. sessilis* showed the opposite patterns. These results suggest that reduced competitive intensity among kin may contribute to the invasion success of *A. philoxeroides*.

As related individuals (i.e. kin) have the same resource requirements, resulting in high niche overlap, they might compete more intensely with each other. However, kin selection theory predicts that individuals compete less intensely with relatives than with strangers, as this would result in higher inclusive fitness for the kin (Hamilton, 1964; Ehlers & Bilde, 2019). In this study, we found that the invasive species indeed exhibited a lower negative relative neighbor effect (RNE) in kin groups than nonkin groups (Figure 3), and that the kin groups produced more biomass than nonkin groups. Therefore, the results for the invasive species support the predictions of kin selection theory. Alternatively, these results could be explained by size-asymmetric competition (Jensen’s inequality), in which greater size asymmetry among nonkin than among kin can result in reduced biomass production of nonkin (Simonsen et al., 2014; Ehlers & Bilde, 2019). However, this is likely not be the case in our study, because—in contrast to the native species—the coefficients of variation (CV) in aboveground biomass of the three individuals per pot did not differ between the kinship treatments of the invasive species (Table S5; Figure S2).

In contrast to the invasive species, the native species had a lower biomass production and the central plants experienced a more negative RNE (i.e. a higher competitive intensity) in the kin treatment than in the nonkin treatment (Table 1; Figure 2a). This is in line with niche-partitioning theory, which predicts that when related individuals compete their fitness will be lower than when unrelated individuals compete (Adler et al., 2018; Chesson, 2018). This is because relatives are phenotypically similar and thus compete more intensely for the same resources than unrelated individuals (Chesson, 2018).As the native species *A. sessilis* exhibits high genetic variation within populations (Geng et al., 2006), it experiences less kin competition, and therefore might—in contrast to the invasive *A. philoxeroides—* not have evolved mechanisms to reduce kin competition.

Phenotypic plastic responses in aboveground architecture may be a mechanism to reduce or avoid negative kin interactions (Ninkovic, 2003; Biedrzycki et al., 2010). In our study, aboveground architecture of central plants did not depend significantly on whether the neighbor plants were kin or nonkin (Table 3, S4, Figure 4). However, in the native species, the neighbor plants produced more branches but less aboveground biomass in the kin treatment than in the nonkin treatment, whereas the opposite was true for the invasive species (Table 4, S3; Figure 5, S1). Since branch distribution is closely correlated with leaf distribution, a greater number of branches can lead to increased shading of adjacent plants and thus intensified competition for light (Pearcy et al., 2005). Thus, our findings suggest that the invasive species altered its aboveground architecture through phenotypic plasticity to reduce kin competition, whereas the native species altered it to intensify kin competition.

In addition to aboveground plastic responses, the outcome of kin interactions has also been linked to shifts in belowground traits such as a reduced root-to-shoot ratio among kin, suggesting less kin competition belowground (Dudley & File, 2007; Murphy & Dudley, 2009; Chen et al., 2012). In our study, the invasive species *A. philoxeroides* showed a weaker relative neighbor effect (RNE) of central plants in the kin treatment and produced higher root biomass overall. However, it did not exhibit significant differences in root allocation between kin and nonkin groups (Table1, S6; Figure 2, S3). Previous studies have shown that the invasive species *A. philoxeroides* improves resource uptake ability through increased root growth, which is crucial for its successful invasion (Wang et al., 2016; Shen et al., 2022). Also, research on other species demonstrated that increased root biomass may reflect soil space exploration rather than competitive response to neighbors (Reynolds & D’Antonio, 1996; Ren et al., 2019). Therefore, the higher root biomass observed in the invasive species likely reflects its inherent resource-acquisition strategy rather than a competitive response to neighbors or kin recognition. It should be noted that roots were measured at the group level in this study, which limits individual-level inferences. Future work is needed to clarify whether and how root allocation responds to neighbor kinship at the individual plant level.

With the intensification of climate change, drought events have become more frequent globally and may impact plant invasions (Sanders et al., 2025). Therefore, our study tested whether drought stress can influence kin interactions. Previous studies have demonstrated that kin interactions can shift from competition to facilitation in stressful environments (He et al., 2013; West et al., 2021). In our experiment, drought stress exerted a stronger negative effect on native species than on invasive species. However, we did not find that drought altered kin or nonkin interactions in either species, which may indicate that the outcomes of kin interactions are relatively robust to environmental variation in the studied system. Alternatively, the drought stress might not have been severe enough to induce a shift in interaction patterns, particularly for the invasive species, which exhibits inherent drought tolerance (Pan et al., 2007).

While central plants of the invasive species showed a much weaker RNE with kin neighbors than nonkin neighbors, they showed weaker kin–nonkin differentiation in aboveground biomass than was the case for the native species. This suggests that the weaker interactions between kin than among nonkin of the invasive species are not due to kin avoidance responses. One possible explanation for the observed pattern is that the overall biomass of the invasive species was much lower than that of the native species, and that significant belowground responses were observed for the invasive plants. It is thus possible that the belowground effects had not yet fully translated into aboveground biomass differences by the time of harvest. Another explanation could be that the weak kin response may be attributed to the generally low genotypic variation reported across Chinese populations of *A. philoxeroides* (Ye et al., 2003; Xu et al., 2003; Wang et al., 2024). Future studies should assess to which degree the designated “kin” and “nonkin” groups of the invasive species were genetically distinct. However, the clear difference in biomass production between the kin and nonkin treatments indicates that the populations of the invasive species were genetically distinct.

## Conclusion

Our study revealed distinct intraspecific interactions with kin and nonkin plants between the invasive *A. philoxeroides* and its native congener *A. sessilis*. By analyzing biomass production and phenotypic plasticity in branching at both the group and individual levels, we provide multiple lines of evidence that the invasive species exhibits weaker competition among kin than among nonkin, whereas the oppositive is true for the native species. This reduced kin competition of the invasive species likely enhances individual growth and population fitness, and may have promoted its invasion success.

## Data accessibility

Should the manuscript be accepted, the data supporting the results will be archived in Figshare, and the data DOI will be included in the article.

## Acknowledgments

We thank Lichao Wang, Huifei Jin, Qingfen Yang and Xue Zhang for help with the plant harvest. YL was supported by the Postdoctoral International Exchange Program for Incoming Postdoctoral Students (YJ20200186).

## Author contributions

XLX and YL conceived the idea and designed the experiment. YL and ZYT performed the experiment. YL and ZYT analyzed the data and wrote the first draft of the manuscript, with further inputs from XLX and MVK.

