## Appendix S1 for "Differential kin interactions between invasive and native plants: evidence from *Alternanthera philoxeroides* and its native congener"

**Table S1** Information on the native and invasive study species.

| Species | Status | Population code | Geographic Area (Province/Company, Country) |
|---|---|---|---|
| *Alternanthera sessilis* | native | pop1 | Hubei, China |
| | | pop2 | Guangdong, China |
| | | pop3 | Jiangxi, China |
| | | pop4 | Guangxi, China |
| *Alternanthera philoxeroides* | invasive | pop1 | Anhui, China |
| | | pop2 | Jiangsu, China |
| | | pop3 | Zhejiang, China |
| | | pop4 | Shanghai, China |

**Table S2** Dummy variable coding for the neighbor treatments. Note that a model including all three dummy variables is equivalent to a model including a single factor with four levels identifying the four different neighbor types.

| Neighbor type | Dummy variable | | |
|---|---|---|---|
| | C1 | C2 | C3 |
| Without neighbor | 1 | 1 | 1 |
| Kin neighbors | 0 | 0 | 1 |
| Mixed neighbors | 0 | 1 | 1 |
| Nonkin neighbors | 0 | 0 | 0 |

**Table S3** Results of linear mixed-effect models used to test the effects of species status (invasive vs. native), watering treatment (wet vs. drought), neighbor type (coded as dummy variables: C1, C2, C3; dummy variables were coded to allow testing of the specific contrasts listed in Figure 1) and all their interactions on aboveground biomass of central plants and neighbor plants per group. Model error distribution, applied transformations, and sample size are indicated in brackets. Significant effects (P < 0.05) are in bold.

| Explanatory variables (fixed and random effects) | df | Aboveground biomass of central plants (Gaussian, log, n=191) | | Aboveground biomass of neighbor plants (Gaussian, log, n=143) | |
|---|---|---|---|---|---|
| | | $\chi2$ | P | $\chi2$ | P |
| Status | 1 | **19.95** | **<0.0001** | **24.82** | **<0.0001** |
| Watering treatment | 1 | **281.05** | **<0.0001** | **276.05** | **<0.0001** |
| C1 (alone vs. with neighbors) | 1 | **89.14** | **<0.0001** | - | - |
| C2 (with mixed neighbors vs. averaged across with kin neighbors and with nonkin neighbors) | 1 | **4.03** | **0.04** | 0.06 | 0.81 |
| C3 (with kin neighbors vs. with nonkin neighbors) | 1 | 0.05 | 0.82 | 1.03 | 0.31 |
| Status × Watering treatment | 1 | 0.00 | 0.97 | 0.11 | 0.74 |
| Status × C1 | 1 | **11.29** | **<0.001** | - | - |
| Status × C2 | 1 | 0.02 | 0.90 | 0.41 | 0.52 |
| Status × C3 | 1 | **5.88** | **0.02** | *3.34* | *0.07* |
| Watering treatment × C1 | 1 | **3.96** | **0.05** | - | - |
| Watering treatment × C2 | 1 | 1.68 | 0.20 | 0.00 | 0.94 |
| Watering treatment × C3 | 1 | 0.54 | 0.46 | 1.95 | 0.16 |
| Status × Watering treatment × C1 | 1 | 0.97 | 0.32 | - | - |
| Status × Watering treatment × C2 | 1 | 0.02 | 0.88 | 2.13 | 0.14 |
| Status × Watering treatment × C3 | 1 | 1.01 | 0.31 | 0.21 | 0.65 |
| **Random** | | SD | | SD | |
| Population | 1 | 0.09 | | 0.04 | |
| Residual | 1 | 0.35 | | 0.28 | |

**Table S4** Results of linear mixed-effect models used to test the effects of species status (invasive vs. native), watering treatment (wet vs. drought), neighbor type (coded as dummy variables:C1 C2, C3; dummy variables were coded to allow testing of the specific contrasts listed in Figure 1) and all their interactions on the log response ratio (LRR) of node number and branch number of central plants. Model error distribution and sample size are indicated in brackets. Significant effects ($P < 0.05$) are in bold.

| Explanatory variables (fixed and random effects) | df | LRR for node number (Gaussian, n=190) | | LRR for branch number (Gaussian, n=190) | |
|---|---|---|---|---|---|
| | | $\chi2$ | P | $\chi2$ | P |
| Status | 1 | 0.50 | 0.48 | 0.18 | 0.67 |
| Watering treatment | 1 | 0.05 | 0.82 | 0.23 | 0.63 |
| C1(alone vs. with neighbors) | 1 | 0.13 | 0.72 | 0.08 | 0.78 |
| C2 (with mixed neighbors vs. averaged across with kin neighbors and nonkin neighbors) | 1 | 2.32 | 0.13 | 2.59 | 0.11 |
| C3(with kin neighbors vs. with nonkin neighbors) | 1 | 0.75 | 0.39 | 0.55 | 0.46 |
| Status × Watering treatment | 1 | 0.00 | 0.97 | 0.11 | 0.75 |
| Status × C1 | 1 | **4.56** | **0.03** | **4.53** | **0.03** |
| Status × C2 | 1 | 1.53 | 0.22 | 0.91 | 0.34 |
| Status × C3 | 1 | 0.00 | 0.97 | 0.00 | 0.96 |
| Watering treatment × C1 | 1 | 0.00 | 0.99 | 0.03 | 0.87 |
| Watering treatment × C2 | 1 | 0.45 | 0.50 | 0.10 | 0.75 |
| Watering treatment × C3 | 1 | 0.01 | 0.93 | 0.02 | 0.88 |
| Status × Watering treatment × C1 | 1 | 2.64 | 0.10 | 2.00 | 0.16 |
| Status × Watering treatment × C2 | 1 | 2.09 | 0.15 | 1.18 | 0.28 |
| Status × Watering treatment × C3 | 1 | 0.00 | 0.95 | 0.06 | 0.81 |
| **Random** | | SD | | SD | |
| Population | 1 | 0.22 | | 0.00 | |
| Residual | 1 | 2.60 | | 2.30 | |

**Table S5** Results of linear mixed-effect models used to test the effects of watering treatment (wet vs. drought), neighbor type (coded as dummy variables: C2, C3; dummy variables were coded to allow testing of the specific contrasts listed in Figure 1) and all their interactions on the coefficients of variation (CV) of aboveground biomass among the three plants in each pot in native and invasive species at mature stage. Model error distribution, applied transformations, and sample size are indicated in brackets. Significant effects (P < 0.05) are in bold.

| Explanatory variables (fixed and random effects) | df | CV of aboveground biomass in native species (Gaussian, log, n=72) | | CV of aboveground biomass in invasive species (Gaussian, log, n=71) | |
|---|---|---|---|---|---|
| | | $\chi2$ | P | $\chi2$ | P |
| Watering treatment | 1 | 1.76 | 0.18 | 0.98 | 0.32 |
| C2 (with mixed neighbors vs. averaged across with kin neighbors and nonkin neighbors) | 1 | 0.13 | 0.72 | 2.66 | 0.10 |
| C3 (with nonkin neighbors vs. with kin neighbors) | 1 | **6.50** | **0.01** | 0.79 | 0.37 |
| Watering treatment × C2 | 1 | 0.95 | 0.33 | 1.19 | 0.27 |
| Watering treatment × C3 | 1 | *3.64* | *0.06* | 0.23 | 0.63 |
| **Random** | | SD | | SD | |
| Population | 1 | 0.12 | | 0.00 | |
| Residual | 1 | 0.58 | | 0.66 | |

**Table S6** Results of linear mixed-effect models used to test the effects of species status (invasive vs. native), watering treatment (wet vs. drought), neighbor type (coded as dummy variables: C1, C2, C3; dummy variables were coded to allow testing of the specific contrasts listed in Figure 1) and all their interactions on root allocation by groups. Model error distribution, applied transformations, and sample size are indicated in brackets. Significant effects (P < 0.05) are in bold.

| Explanatory variables (fixed and random effects) | df | Root allocation (Gaussian, log, n=191) | |
|---|---|---|---|
| | | $\chi 2$ | P |
| Status | 1 | **1.77** | **0.18** |
| Watering treatment | 1 | **34.80** | **<0.001** |
| C1 (alone vs. with neighbors) | 1 | **13.71** | **<0.001** |
| C2 (with mixed neighbors vs. averaged across with kin neighbors and nonkin neighbors) | 1 | 0.29 | 0.59 |
| C3 (with kin neighbors vs. with nonkin neighbors) | 1 | 0.42 | 0.52 |
| Status × Watering treatment | 1 | 0.65 | 0.42 |
| Status × C1 | 1 | *2.88* | *0.09* |
| Status × C2 | 1 | *3.38* | *0.07* |
| Status × C3 | 1 | 0.48 | 0.49 |
| Watering treatment × C1 | 1 | 2.00 | 0.16 |
| Watering treatment × C2 | 1 | 0.21 | 0.65 |
| Watering treatment × C3 | 1 | 0.09 | 0.77 |
| Status × Watering treatment × C1 | 1 | 0.05 | 0.82 |
| Status × Watering treatment × C2 | 1 | 2.18 | 0.14 |
| Status × Watering treatment × C3 | 1 | 0.08 | 0.77 |
| **Random** | | SD | |
| Population | 1 | 0.14 | |
| Residual | 1 | 0.28 | |
