## Appendix S2 for "Differential kin interactions between invasive and native plants: evidence from *Alternanthera philoxeroides* and its native congener"

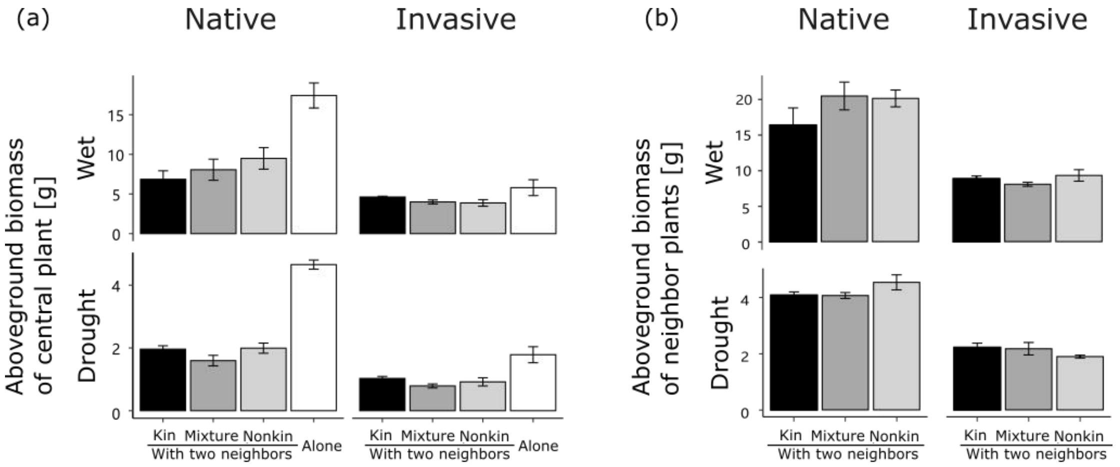


**Figure S1 Aboveground biomass of central plants (a) and neighbor plants (b) in treatments with kin neighbors (black), with mixed neighbors (dark grey), with nonkin neighbors (light grey) and without neighbors (alone; white) for the native and invasive species under wet and drought conditions.** Error bars are standard errors of the mean of population-means (n = 4 populations; each population mean was based on 3 individual plants).

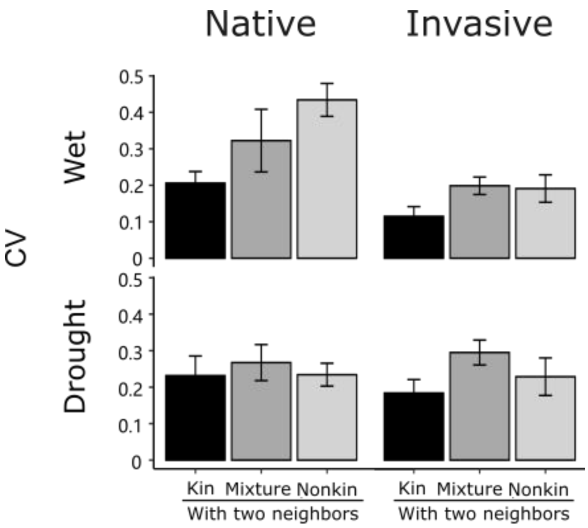


**Figure S2 Coefficient of variation (CV) of aboveground biomass among the three plants within each pot.** Plants were grown in kin groups (black), mixed groups (dark grey), or nonkin groups (light grey) for the native and invasive species under wet and drought conditions. Error bars are standard errors of the mean of population-means (n = 4 populations; each population mean was based on 3 pots, each with three individual plants).

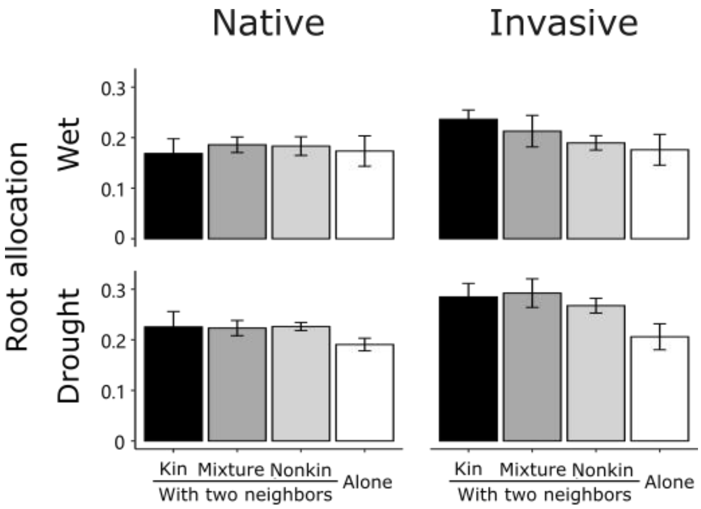


**Figure S3 Root allocation (root-shoot ratio) for the native and invasive species under wet and drought conditions.** Plants were grown in kin groups (black), mixed groups (dark grey), nonkin groups (light grey), or alone (white). Error bars are standard errors of the mean of population-means (n = 4 populations; each population mean was based on 3 pots).
